# Quantitative Temporal Analysis of Protein Dynamics in Maladaptive Cardiac Remodeling

**DOI:** 10.1101/270959

**Authors:** Daniel B. McClatchy, Yuanhui Ma, David A. Liem, Dominic C.M. Ng, Peipei Ping, John R. Yates

**Author notes:** Co-authors (Daniel B. McClatchy, Yuanhui Ma).

## Abstract

Maladaptive cardiac remodeling (MCR) is a complex dynamic process common to many heart diseases. MCR is characterized as a temporal progression of global adaptive and maladaptive perturbations. The complex nature of this process clouds a comprehensive understanding of MCR, but greater insight into the processes and mechanisms has the potential to identify new therapeutic targets. To provide a deeper understanding of this important cardiac process, we applied a new proteomic technique, PALM (Pulse Azidohomoalanine in Mammals), to quantitate the newly-synthesized protein (NSP) changes during the progression of isoproterenol (ISO)-induced MCR in the mouse left ventricle. This analysis revealed a complex combination of adaptive and maladaptive alterations at acute and prolonged time points including the identification of proteins not previously associated with MCR. We also combined the PALM dataset with our published protein turnover rate dataset to identify putative biochemical mechanisms underlying MCR. The novel integration of analyzing NSPs together with their protein turnover rates demonstrated that alterations in specific biological pathways (e.g., inflammation and oxidative stress) are produced by differential regulation of protein synthesis and degradation.

## Introduction

Heart failure (HF) and maladaptive cardiac remodeling (MCR) are both common stages of many heart diseases with increasing incidence and prevalence in the United States, posing major public health problems (McMurray JJ et al, Circulation 2010: 111, 1233-1241). MCR is a multifactorial disease and results not only from cardiac overload or injury but also from a complex interplay among genetic, inflammatory, biochemical, and neuro-hormonal alterations (Shimizu and Minamino 2016). Accordingly, the MCR process involves prolonged over-stimulation of the sympathetic nervous system (SNS) followed by the release of catecholamines to increase both the heart rate and individual cellular contractility, thereby creating a compensatory adaptive response that transiently normalizes the biomechanical stress and optimizes cardiac contractility. Catecholamines are crucial stress hormones and regulators of heart function via stimulation of beta-adrenergic receptors (β-ARs) (Guggliam et al, JMCC 2013 Apri;57:47-58). Chronic over-stimulation, however, augments the MCR process, eventually leading to cardiac dysfunction and HF (Bristow, Ginsburg et al. 1982, Lohse, Engelhardt et al. 2003). In the early stages MCR exhibits thickening of the ventricular and interventricular walls and is characterized by increasing size of the cardiomyocytes, changes in the organization of the sarcomeric structure, and increased protein synthesis (Drews, Tsukamoto et al. 2010). Similarly, persistent pharmacological β-AR stimulation with the agonist isoproterenol (ISO) can cause left ventricle hypertrophy and HF in laboratory animals (Rona, Chappel et al. 1959, Taylor and Tang 1984). Numerous studies have suggested that chronic ISO application stimulates structural and metabolic remodeling, which alters fatty acid utilization, glucose homeostasis, and mitochondrial function (Colucci 1997, van Bilsen, Smeets et al. 2004). To advance our understanding of the underlying molecular mechanisms of MCR and HF, it is important to identify how proteome alterations gradually manifest during these pathophysiological stages over time. Identification of proteins that drive MCR may lead to novel therapeutic drug targets or clinical biomarkers for cardiac pathologies.

Quantitative proteomics has great potential to better understand the myriad of proteome alterations during the progression of ISO-induced MCR. We recently developed the quantitative proteomic technique PALM (Pulse Azidohomoalanine Labeling in Mammals) to improve the temporal resolution of quantitative proteomics (McClatchy, Ma et al. 2015). PALM quantitates newly-synthesized proteins (NSPs). Since the NSP sub-proteome in theory should be the first to respond to perturbations, quantitation of NSPs should be more sensitive than quantitation of the whole proteome. PALM relies on azidohomoalanine (AHA), which is a non-canonical amino acid that is accepted by the endogenous methionine tRNA and inserted into proteins in vivo (Dieterich, Link et al. 2006). AHA is inserted into the mouse proteome through a customized AHA diet. The AHA diet is given to mice within a discrete time period to label NSPs in response to a perturbation. Proteins incorporating AHA can be covalently linked in vitro to a biotin-alkyne with click chemistry. Through standard biotin enrichment strategies, the static or “old” proteome is removed, and the remaining NSPs can be identified and quantitated using mass spectrometry (MS). In this study, we applied the PALM strategy to the ISO-induced MCR mouse model, to enhance our mechanistic understanding of cardiac proteome remodeling in this pathophysiological process. We quantified newly synthesized proteins through an acute phase, during which the MCR process is at its highest rate, and a prolonged phase, during which the MCR process has reached a plateau. In the normal heart, homeostasis of continuous protein synthesis and degradation is essential to maintain cardiac function at the cellular and whole-organ level (Lam, Wang et al. 2014). Normal protein homeostasis through the protein turnover cycle is altered during MCR and HF through factors such as hypertrophic signaling (Shimizu and Minamino 2016), calcium regulation (Goonasekera, Hammer et al. 2012), inflammatory reactions (Shimizu and Minamino 2016), and oxidative stress (Goirdano FJ et al, JCI 2005;115(3);500-508). For the first time, this study integrates two novel datasets using previously published protein turnover rates in the ISO-induced MCR mouse model with the NSP quantitation in response to ISO treatment. This integrated analysis provides unique insight into the total balance of protein synthesis and degradation in two compared groups (i.e., sham group vs. ISO group) and highlights the mechanisms that underlie proteomic changes in MCR. It also establishes a novel experimental and bioinformatic approach that is generally applicable to animal models of disease.

## Methods

### Animals/Surgery/Tissue Collection

MCR was induced by continuous ISO treatment for 14 days in male C57/BL6 wildtype mice (aged 8-12 weeks) using microosmotic pumps (Alzet, Model 1002). The microosmotic pumps were implanted subcutaneously in the posterior neck area of the mouse under 2% isoflurane (vaporized in oxygen) anesthesia. One group of mice received continuous treatment of ISO with a dose of 15 mg/kg/day for a total period of 14 days. As a control, a second group of mice underwent sham treatment by vehicle (PBS) (**Figure 1A**).

**Figure 1:**
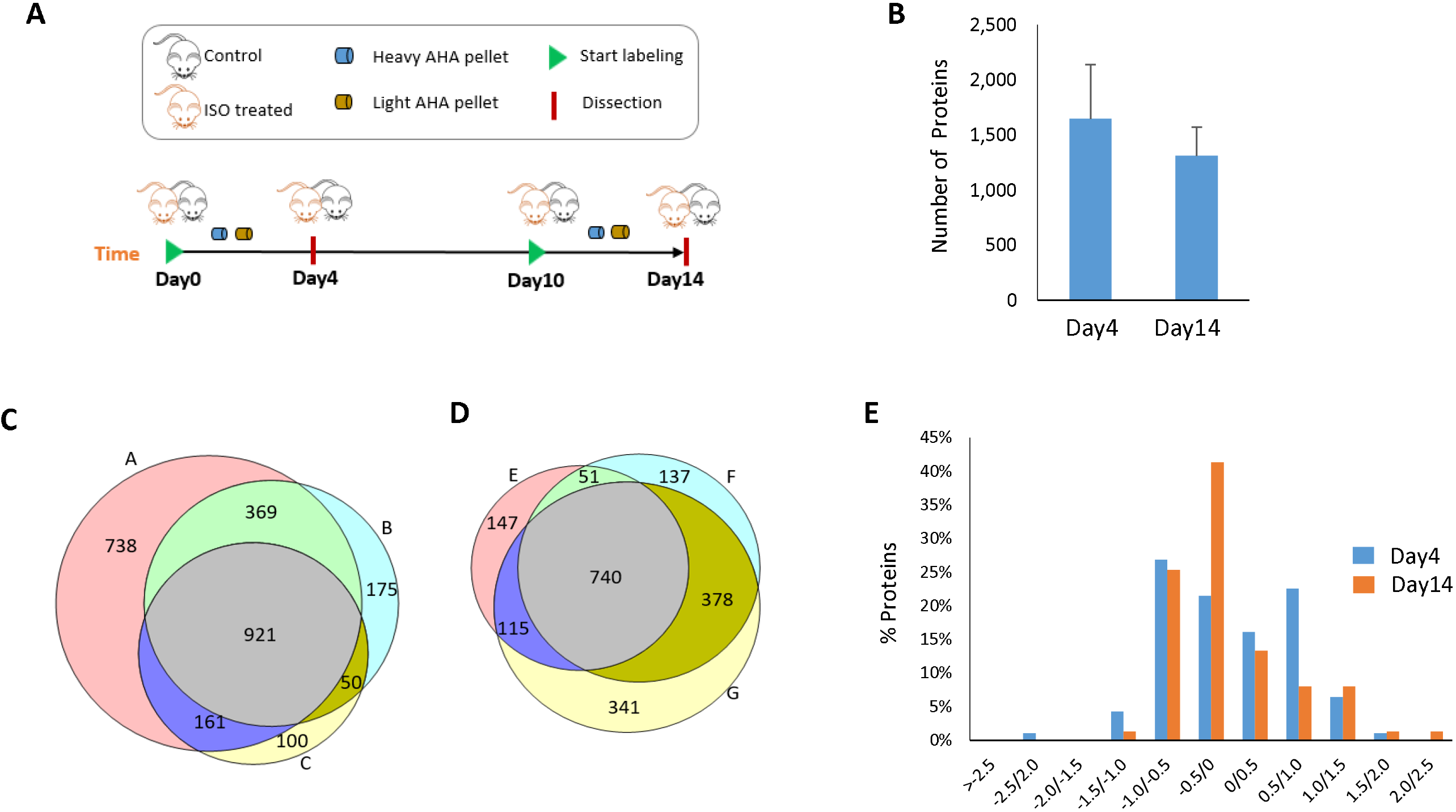
Protein identification of Day 4 and Day 14 time points. **A)** Schematic workflow. Surgically implanted osmotic pumps delivered ISO (15 mg/kg/d) or saline (i.e., control) to mice for 14 days. The mice were fed a diet of either light AHA (L-AHA) or heavy AHA (H-AHA) for four days at different time points. Left ventricles (LVs) were dissected after labeling. **B)** The number of NSPs identified at Day 4 and Day 14 are similar. Average protein numbers of three replicates for each time point were plotted. Error bar represents standard deviation. The overlap of protein identifications of three biological replicates are from **(C)** Day 4 (replicate A, B, and C) analysis and **(D)** Day 14 (replicate E, F, and G) analysis. **E)** Significant NSPs (p< 0.05 using a one sample t-test) were plotted in a histogram with their fold change (ISO/control) on the x-axis. There were 93 and 75 proteins that were significantly different between the ISO and control hearts in Day 4 and Day 14, respectively.

AHA-Heavy pellets were fed to the ISO group and AHA-Light pellets to the control group. For efficient labeling as well as to study two different phases in the MCR process, we collected samples on Day 4 and Day 14 in both the ISO group and the control group. To collect samples on Day 4, we commenced AHA labeling on Day 0. To collect samples on Day 14, we commenced AHA labeling on Day 10. To conduct our AHA labeling analyses after the 14-day period treatment, whole heart, skeletal muscle, liver, kidney, and brain were collected after sacrificing the mice. In contrast to sham treatment, the heart weight-body weight ratios (HW/BW) were increased in the ISO group on Day 4 and Day 14, confirming cardiac remodeling. Cardiac tissues were collected from left ventricle (LV) and right ventricle (RV) and rinsed in PBS.

### Tissue homogenization

LVs were transferred to Precellys CK14 tubes containing 1.4 mm ceramic beads, and 0.5mm disruption beads (Research Products International Corp.) and 500ul PBS were added. The homogenization was performed on Bertin Precellys bead-beating system: 6500 rpm, 3 times, 20 seconds internal. Then 0.5% SDS was added and the homogenates were sonicated for 30 secs. A BCA protein assay was performed on each sample.

### Click chemistry

2.5mg protein of LV from heavy AHA (H-AHA)-labeled ISO mice or sham mice and 2.5mg protein of LV from light AHA (L-AHA)-labeled sham mice or ISO mice were mixed together as one biological replicate. For each biological replicate, the H-AHA/L-AHA mixture was divided into twenty aliquots (0.25 mg/aliquot). A click reaction was performed on each aliquot as previously published (Speers and Cravatt 2009). Briefly, for each click reaction the following reagents were added in this order: 1) 30 ul of 1.7mM TBTA, 2) 8 ul of 50mM Copper Sulfate, 3) 8 ul of 5mM Biotin-Alkyne (C_21_H_35_N_3_O_6_S, Click Chemistry Tools), and 4) 8 ul of 50mM TCEP. PBS was then added to a final volume of 400 ul and incubated for 1 hour at room temperature. Methanol/Chloroform precipitation was performed, and the precipitated protein was combined so that there would only be one pellet per 5mg starting material.

### Digestion and biotin peptide enrichment

Biotin peptide enrichment was performed as previously described (Schiapparelli, McClatchy et al. 2014). Briefly, precipitated pellets were resuspended in 100ul 8M urea and 100ul 0.2% MS-compatible surfactant ProteaseMAX (Promega) in 50mM ammonium bicarbonate, then reduced, alkylated, and digested with trypsin as previously described (Schiapparelli, McClatchy et al. 2014). The digestion was then centrifuged at 13,000 × g for 10 min. The supernatant was transferred to a new tube, and the pellet was resuspended with PBS and centrifuged at 13,000 × g for 10 min. Supernatants were combined, and 150 ul of neutravidin agarose resin (Thermo Scientific) was added. The resin was incubated with the peptides for 2 hours at room temperature while rotating. The resin was then washed with 1ml PBS, PBS with 5% acetonitrile, PBS, and a final wash of distilled water. The peptides were eluted from the resin twice with 150ul 80% acetonitrile, 0.2% formic acid, and 0.1% TFA on shaker for 5 min at room temperature, and another two times on a shaker at 70 °C. All elutions were transferred to a single new tube. Prior to MS analysis, the samples were dried in a speed-vac, and dried peptides were resolubilized in buffer A (5% ACN, 95% water, 0.1% formic acid).

### Mass spectrometry

Soluble peptides were pressure-loaded onto a 250-μm i.d. capillary with a kasil frit containing 2.5 cm of 10-μm Jupiter C18-A material (Phenomenex) followed by 2.5 cm of 5-μm Partisphere strong cation exchanger (Whatman). This column was washed with buffer A after loading. A 100-μm i.d. capillary with a 5-μm pulled tip packed with 15 cm of 4-μm Jupiter C18 material (Phenomenex) was attached to the loading column with a union, and the entire split-column (loading column–union–analytical column) was placed in line with an Agilent 1100 quaternary HPLC (Palo Alto). The sample was analyzed using MudPIT, which is a modified 11-step separation described previously (Washburn, Wolters et al. 2001). The buffer solutions used were: buffer A, buffer B (80% acetonitrile/0.1% formic acid), and buffer C (500-mM ammonium acetate/5% acetonitrile/0.1% formic acid). Step 1 consisted of a 10 min gradient from 0-10% buffer B, a 50 min gradient from 10-50% buffer B, a 10 min gradient from 50-100% buffer B, and 20 min from 100% buffer A. Steps 2 consisted of 1 min of 100% buffer A, 4 min of 20% buffer C, a 5-min gradient from 0-10% buffer B, an 80-min gradient from 10-45% buffer B, a 10-min gradient from 45-100% buffer B, and 10 min of 100% buffer A. Steps 3-9 had the following profile: 1 min of 100% buffer A, 4 min of X% buffer C, a 5-min gradient from 0-15% buffer B, a 90-min gradient from 15-45% buffer B, and 10 min of 100% buffer A. The buffer C percentages (X) were 30, 40, 50, 60, 70, 80, and 100%, respectively, for the following 7-step analysis. In the final two steps, the gradient contained: 1 min of 100% buffer A, 4 min of 90% buffer C plus 10% B, a 5-min gradient from 0-10% buffer B, an 80-min gradient from 10-45% buffer B, a 10-min gradient from 45-100% buffer B, and 10 min of 100% buffer A. As peptides eluted from the microcapillary column, they were electrosprayed directly into an Elite mass spectrometer (ThermoFisher) with the application of a distal 2.4-kV spray voltage. A cycle of one full-scan FT mass spectrum (300-1600 m/z) at 240,000 resolution, followed by 20 data-dependent IT MS/MS spectra at a 35% normalized collision energy, was repeated continuously throughout each step of the multidimensional separation. Application of mass spectrometer scan functions and HPLC solvent gradients were controlled by the Xcalibur data system.

### Data analysis

Both MS1 and MS2 (tandem mass spectra) were extracted from the XCalibur data system format (.RAW) into MS1 and MS2 formats using in-house software (RAW_Xtractor)(McDonald, Tabb et al. 2004). MS/MS spectra remaining after filtering were searched with ProLuCID (Xu, Venable et al. 2006) against the UniProt_Mouse_03-25-2014 concatenated to a decoy database, in which the sequence for each entry in the original database was reversed (Peng, Elias et al. 2003). All searches were parallelized and performed on a Beowulf computer cluster consisting of 100 1.2-GHz Athlon CPUs (Sadygov, Eng et al. 2002). No enzyme specificity was considered for any search. The following modifications were searched for analysis: a static modification of 57.02146 on cysteine for all analyses, a differential modification of 452.2376 on methionine for AHA-biotin-alkyne, or 458.2452 for H-AHA-biotin-alkyne. ProLuCID results were assembled and filtered using the DTASelect (version 2.0) program (Tabb, McDonald et al. 2002, Cociorva, D et al. 2007). DTASelect 2.0 uses linear discriminant analysis to dynamically set XCorr and DeltaCN thresholds for the entire dataset to achieve a user-specified false discovery rate (FDR). In DTASelect, the modified peptides were required to be fully tryptic, less than 5-ppm deviation from peptide match, and a FDR at the spectra level of 0.01. The FDRs are estimated by the program from the number and quality of spectral matches to the decoy database. For all datasets, the protein FDR was < 1% and the peptide FDR was < 0.5%. The MS data was quantified (i.e., generate heavy/light ratios) using the software pQuant(Liu, Song et al. 2014), which uses the DTASelect and MS1 files as the input. pQuant assigns a confidence score to each heavy/light ratio from zero to one. Zero, the highest confidence, means there is no interference signal, and one means the peptide signals are almost inundated by interference signals (i.e., very noisy). For this analysis, only peptide ratios with sigma less than 0.1 and intensity over 1e^5^ were allowed for quantification.

### Pathway analysis

The significantly changed proteins, combined with unquantifiable large changes, were listed as input for functional pathway analysis performed by the software Ingenuity Pathway Analysis (IPA) (Calvano, Xiao et al. 2005). In addition, the significantly changed proteins combined with unquantifiable large changes of Day 4 and Day 14 were analyzed by STRING (Szklarczyk, Franceschini et al. 2015) using high confidence score settings and visualized by Cytoscape. Cell localization information was obtained from Uniprot (http://www.uniprot.org/).

### Statistical analysis

One sample t-test was employed to determine the significantly altered NSP at Day-4 ISO vs. Day 4control and Day 14 ISO vs. Day 14 control. Significantly enriched pathways were determined by IPA. In **Figure 4B**, one-way ANOVA was performed, followed by Bonferroni’s post-hoc test, using the software Prism.

## Results

Surgically implanted osmotic pumps delivered ISO (15 mg/kg/d) or PBS (i.e., control) to mice for 14 days (**Figure 1A**). The mice were fed a diet of either light AHA (L-AHA) or heavy AHA (H-AHA) for four days at two different time points. To determine the acute effects of ISO, mice were given the AHA diet during the first four days of ISO delivery (from Day 0 to Day 4), during which the cardiac remodeling process rate was at its maximum. To determine the prolonged effects of ISO, mice were given the AHA diet the last four days of the ISO treatment (from Day 10 to Day 14), during which the cardiac remodeling process was at a plateau. Mice were sacrificed at Day 4 or Day 14, and the LVs were quickly dissected, rinsed with PBS, and snap frozen. The LVs were homogenized, and L-AHA and H-AHA tissues were mixed 1:1 (wt/wt). Day 4 ISO heart tissue was mixed with Day 4 control heart tissue, and Day 14 ISO heart tissue was mixed with Day 14 control heart tissue. Three biological replicates were analyzed for each L-AHA/H-AHA mixture (six mice per time point). Within each time point analysis, a label swap was performed between conditions to remove any possible effects of the heavy label on the proteome. Click chemistry was performed on these mixtures to covalently add a biotin-alkyne. After a tryptic digestion, the AHA peptides were isolated with neutravidin beads. These modified peptides were eluted from the beads and then identified by MS.

### Identification of Newly Synthesized Proteins at Day 4 and Day 14 of Beta-adrenergic-induced Maladaptive Cardiac Remodeling

A similar number of NSPs were identified in the Day 4 and Day 14 analyses (Figure1B **and** SuppTable 1). There were 921 and 740 NSPs identified in all three biological replicates for Day 4 and Day 14 analyses, respectively (Figure 1C **and** 1D), with 608 proteins identified at both time points in all biological replicates. Next, the ion chromatograms for the light and heavy AHA peptide pairs were extracted, and heavy/light ratios were calculated. There were 93 and 75 proteins that were statistically different between the ISO and control hearts in Day 4 and Day 14, respectively (Figure 1E **and SuppTables 2 & 3**). The significant proteins in Day 4 time points were evenly distributed over their fold change (ISO/control), while those in the Day 14 time points were not, with almost 70% of these proteins down-regulated in the ISO group. One caveat to MS quantitative analysis with heavy stable isotopes is that very large changes may prevent a heavy/light ratio from being generated (McClatchy, Liao et al. 2007). When there are very large changes between a light and heavy protein, only one is detected by the mass spectrometer due to its limited dynamic range. Thus, when only the abundance of one peptide (i.e., light or heavy) is measured, a light/heavy ratio cannot be calculated. To identify these unquantifiable large changes, the datasets were searched for proteins identified in all three biological replicates in one condition and not at all in the other (i.e., identified in three Day 4 ISO mice but never identified any of the Day 4 control mice) (**SuppTable 4 & 5**). In the Day 4 acute analysis, 3 proteins were identified in all three ISO mice but not in any control mice, suggesting that these proteins were up-regulated in MCR. Conversely, 9 proteins were identified in all three Day 4 control mice but not in any of the Day 4 ISO mice, suggesting these proteins are down-regulated in MCR. In the Day 14 analysis, 11 proteins were identified in all three ISO mice but not in any control mice, and 1 protein was identified in the control mice and not in the ISO mice.

### Functional Protein Network in Isoproterenol-induced Maladaptive Cardiac Remodeling

All of the proteins identified as unquantifiable large alterations were combined with the statistically significant proteins for further analysis. There were 266 proteins quantified confidently in both the Day 4 and Day 14 analyses (i.e., 6 MS analyses) with a correlation value of r=0.69, suggesting some similar individual protein perturbations between these time points (**Figure 3B**). Eighteen NSPs were statistically significant at both time points, indicating these changes occur during MCR with the highest confidence (Figure 3C **and SuppTable 6**). The protein with the largest change was Ubiquitin carboxyl-terminal hydrolase 13 (USP13), which was identified in all ISO-treated hearts, but never identified in a control heart. These common significantly changed proteins all exhibited the same trend at both time points, except the protein titin (TTN). Next, a functional network for MCR was generated using the STRING database to identify direct and indirect (i.e., functional) interactions within the dataset (**Figure 2**) (Szklarczyk, Franceschini et al. 2015). The proteins were annotated with multiple subcellular compartments, including the nucleus, mitochondria, endoplasmic reticulum, plasma membrane, and the extracellular space. The most striking trend from this network analysis was that 96% of the proteins that localized to the mitochondria were down-regulated in MCR. In contrast, proteins involved in inflammation were all observed to be significantly up-regulated in the MCR network. Next, pathway analysis was performed separately on the acute and prolonged effects of ISO (**Figure 3A**). The altered NSPs were significantly enriched in four pathways at both time points: mitochondrial dysfunction, oxidative phosphorylation, inflammatory response, and X receptor (LXR)/retinoid X receptors (RXR) activation. All of these pathways have previously been reported to be altered in cardiac remodeling (Zhou, Sucov et al. 1995, Epelman, Lavine et al. 2014, Brown, Perry et al. 2017). Ten additional pathways were significantly enriched, but these pathways were only significant at the Day 14 time point.

**Figure 2:**
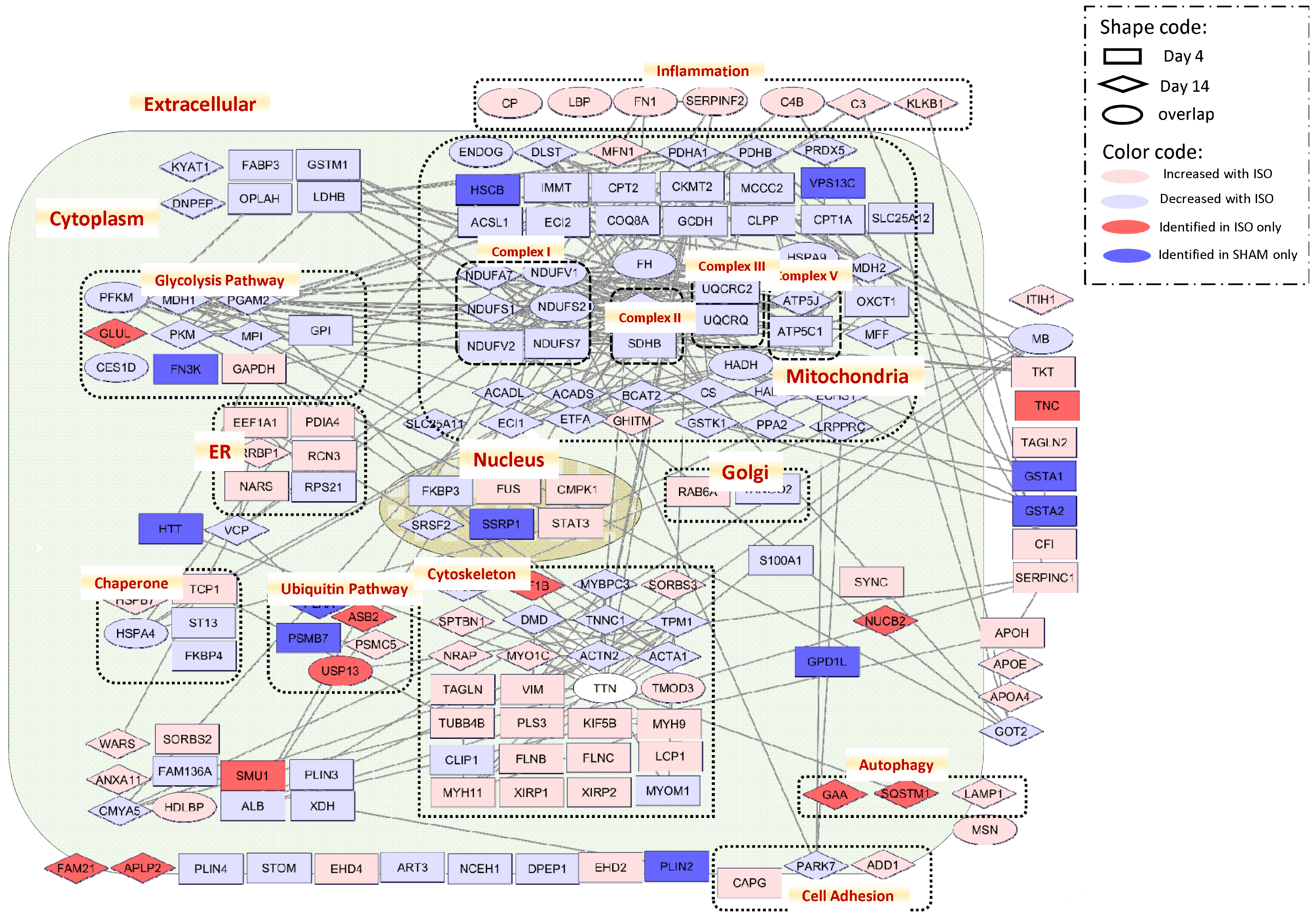
Functional MCR network. The protein network and localization were analyzed by STRING and visualized by Cytoscape. The log ratio used in visualization was the average ratio of replicates. The white ellipse represents the protein Titin (TTN), which was found up-regulated at Day 4 of ISO but down-regulated at Day 14 of ISO.

### Newly Synthesized Proteins in MCR Network associated Cardiac Diseases

After demonstrating that many biological pathways altered in our MCR network have previously been associated with cardiac dysfunction, we next determined if specific proteins in our MCR network have previously been associated with MCR or other cardiac disorders. Twenty-five proteins were found to be directly linked to human cardiac disorders or animal models of cardiac disorders in previous publications (Figure 3D **and SuppTable 7**). Fifty percent of these proteins were significantly altered in both the acute (4 days) and prolonged (14 days) beta-adrenergic stimulation. Mutations in six genes reported to cause inherited human cardiomyopathy (TTN, tropomyosin alpha-1 chain (TPM1); myozenin-2 (MYOZ2), troponin C (TNNC1); myosin-binding protein C (MYBPC3); and alpha-actinin-2 (ACTN2)) were all down-regulated in the MCR network. This is consistent with the human mutations preventing expression or function of these proteins to promote cardiac dysfunction and confirms the relevance of the ISO model to human pathology. The one exception was the contractile protein TTN, which was significantly up-regulated at Day 4 and down-regulated at Day 14. In addition, mutations in two other genes, lysosomal alpha-glucosidase (GAA) and dystrophin (DMD), correlated to Pompe Disease and Duchenne Muscular Dystrophy, respectively, have symptoms that also include cardiac dysfunction and heart failure (Lim, Li et al. 2014, van Westering, Betts et al. 2015). Other proteins in the MCR network (i.e., fatty acid-binding protein (FABP3), fibronectin (FN1), protein deglycase DJ-1 (PARK7)) have been described as altered in or associated with human cardiomyopathy, but a direct genetic cause is not supported. For example, FABP3 is a heart-enriched cytosolic protein responsible for fatty acid transportation. It has been proposed as a useful acute biomarker for human cardiac dysfunction because it is detected in the blood hours after myocardial damage (McLean, Huang et al. 2008). Other proteins (i.e., long-chain-fatty-acid-CoA ligase 1 (ACSL1), ATP-dependent 6-phosphofructokinase (PFKM), fumarate hydratase (FH1)) have been associated with MCR in animals. For example, newly synthesized FH1 was reduced significantly in the MCR network. This protein regulates endogenous levels of fumarate, which is a metabolite of the citric acid cycle and reported to be cardioprotective. Knocking out the FH1 gene in mice increased fumarate levels, which was demonstrated to be cardioprotective against global cardiac ischemia (Ashrafian, Czibik et al. 2012). This correspondence, along with previously published cardiac disease research, suggests great potential to increase the understanding of MCR from the proteins without previous links to cardiac dysfunction.

**Figure 3:**
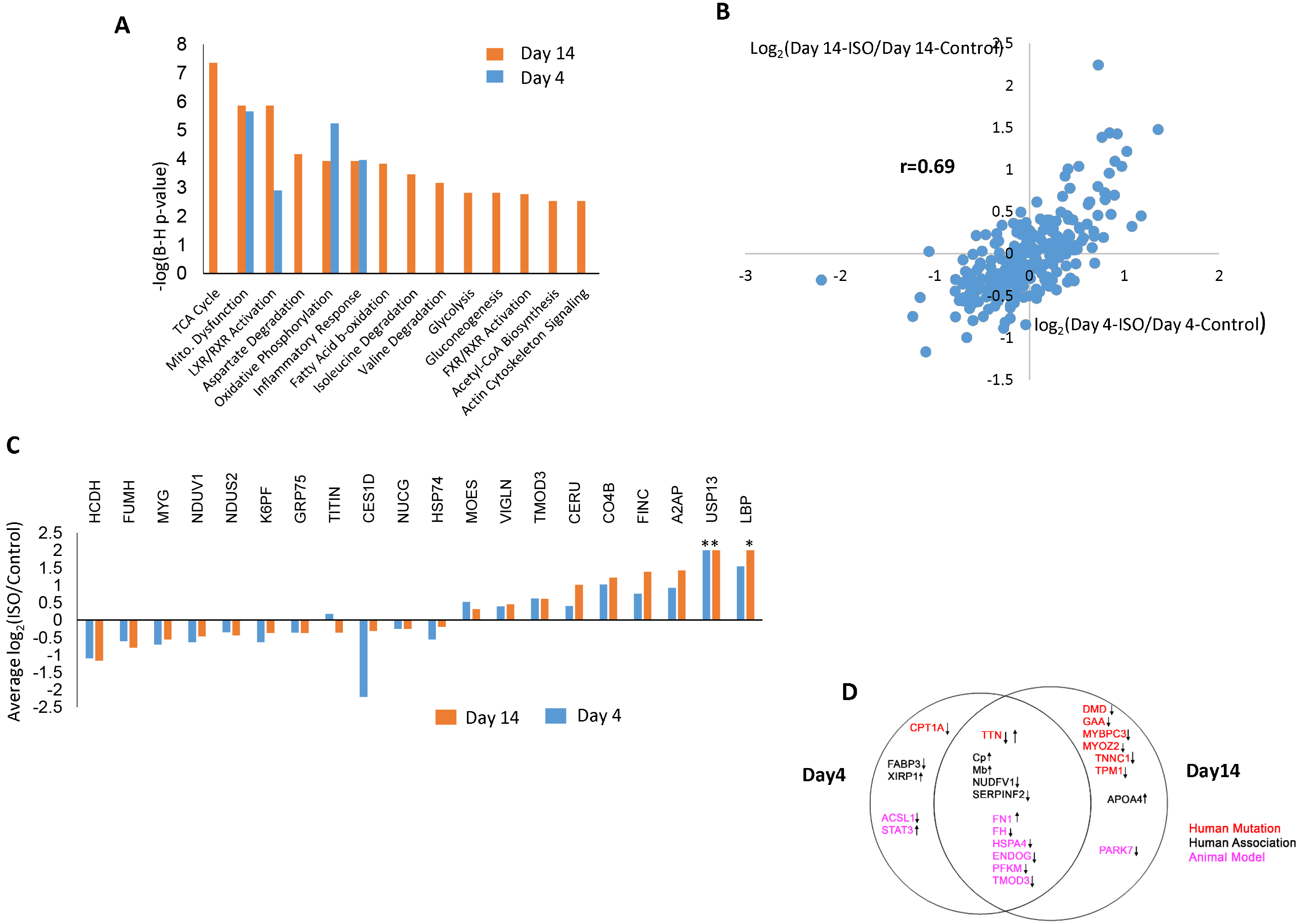
Comparison of acute and chronic ISO treatment. **A)** Pathway analysis on NSPs with significant changes and unquantifiable large changes at acute and chronic time points. **B)** Correlation of NSPs quantified at both acute and chronic analyses. The average of proteins ratios (ISO/control) calculated from 3 biological replicates of each time point were plotted to address correlation between the two datasets. **C)** NSPs quantified at both acute and chronic analyses. “*” represents an unquantifiable large change. **D)** Significantly altered NSPs previously associated with human cardiac diseases or animal models. Manual literature curation associated quantified NSPs in this study with previous reports of human genetic mutations causing cardiac disease (red), biomarker of, or association with, human cardiac disease (black), or association with animal models of cardiac disease (pink). Arrows indicate an up- or down-regulation of the NSP with ISO in this study.

### Integration of Newly Synthesized Proteins and Protein Turnover Rates

To gain insight into how MCR alters cardiac proteome homeostasis, the 14-day dataset was compared with our own published study, which quantified cardiac protein turnover rates using an identical 14 day ISO treatment (Lam, Wang et al. 2014). There were 270 proteins with quantified NSP (ISO/control) ratios and quantified protein turnover (ISO/control) rates (**SuppTable 8**). This comparison dataset was divided into four scenarios: 1) decreased NSP with increased turnover rate; 2) decreased NSP with decreased turnover rate; 3) increased NSP with decreased turnover rate; and 4) increased NSP with increased turnover rate. Changes in protein turnover rates can result from altered rates of degradation or synthesis. Since NSPs are linked directly to protein synthesis, this allowed us to postulate potential mechanisms for each of these observed trends (**Table 1**). Arguably, the simplest scenario for an increase in protein turnover is an increase in protein synthesis, and 30% of proteins in this comparison dataset exhibited this trend. On the other hand, a decrease in protein turnover could result from a decrease in protein synthesis, and this trend was observed in 9% of the proteins. The most frequently observed trend (50%) was a decrease in NSPs with ISO treatment coupled with an increase in turnover rate. We postulate that this correlates to an increase in protein degradation. The last trend, increased NSPs with decreased turnover rate, could represent a decrease in protein degradation but no change in synthesis. The absolute NSP changes were significantly the largest with the trend of a decrease in NSP with ISO treatment coupled with an increase in turnover rate (**Figure 4B**). Importantly, different biological pathways were enriched in each of the trends (**Figure 4A**). The maturation of phagosomes, which remove apoptotic cells, was enriched in proteins with increased NSPs and turnover, indicating that apoptosis of cardiomyocytes is involved in remodeling, as previously reported (Narula, Haider et al. 1996, Olivetti, Abbi et al. 1997). We postulate that an increase in translation is employed to up-regulate this pathway in response to apoptosis. It was observed that a decrease in translation was employed to decrease fatty acid oxidation, which was significantly enriched in proteins with decreased NSPs and decreased turnover. This corresponds to the reported reduction in the protein expression of this pathway in cardiac remodeling as the fuel source shifts from fatty acids to carbohydrates (Akhmedov, Rybin et al. 2015). The trend of decreased NSPs coupled with increased turnover was significantly enriched in mitochondrial signaling, suggesting that mitochondrial proteins have increased rates of degradation in MCR. Increases in NSP due to decreased degradation were enriched in the cardioprotective LXR activation pathway and inflammation. Overall, this analysis suggests that MCR alters different signaling pathways employing different regulatory mechanisms.

**Figure 4:**
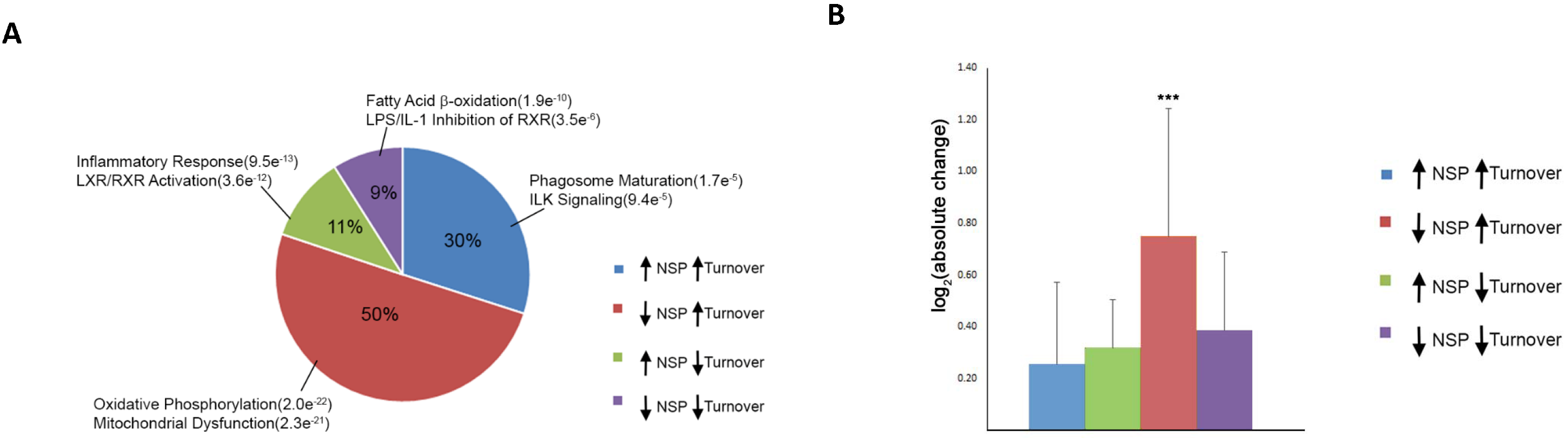
**A)** Percentage of trends identified from the comparison of NSP ratios and protein turnover rates in response to ISO. Each trend is annotated with the top two significantly enriched pathways. The number in parentheses represents the p-value for pathway enrichment. **B)** Comparison of the absolute NSP changes associated with each of the four NSP/turnover trends. One-way ANOVA analysis was performed, and a significant (p < 0.0001) difference was observed between the four trends. A Bonferroni’s Multiple Comparison post-hoc test was then performed, and a significant (***p<0.001) difference was observed in the “down NSP/up turnover” trend and each of the other trends.

## Discussion

The cardiac proteome consists of thousands of proteins interacting in stable and transient complexes as well as numerous signaling pathways that maintain homeostasis and optimal cardiac function. MCR is a process involved in many cardiac disorders and results not only from cardiac overload or injury but also from perturbations in a spectrum of biological processes, eventually leading to cell death (Shimizu and Minamino 2016). To study the temporal progression of alterations that underlie MCR from this incredibly complex proteome, this study sought to reduce proteome complexity by quantifying the NSP sub-proteome using the methionine analog AHA. AHA has been routinely employed to identify and quantify NSPs by MS in cultured cells (Dieterich, Link et al. 2006, Zhang, Bowling et al. 2014). The PALM method was recently reported to incorporate AHA into mouse tissue proteomes during a discrete temporal window (McClatchy, Ma et al. 2015). Taking advantage of the PALM strategy, the acute (i.e., Day 4) and prolonged (i.e., Day 14) proteomic changes were quantified in an ISO-induced mouse model of MCR. Thousands of NSPs were identified and quantified from the LV, revealing temporal changes in MCR. Accordingly, the proteins that were significantly altered between ISO and control hearts were assembled to form a temporal MCR network. In correspondence with published studies on MCR, a large variety of different protein functions were altered from multiple subcellular compartments. Nevertheless, many proteins within our MCR network are connected by direct or functional interactions.

A unique aspect of our MCR network is the temporal dynamics. MCR involves a transition from adaptive or cardioprotective to maladaptive changes. One may hypothesize that the adaptive changes may be present in the acute phase while the maladaptive manifest in the prolonged phase. There is some evidence to support this hypothesis in our MCR network. Six mutated genes known to cause inherited human cardiomyopathy were quantified as down-regulated by ISO in our MCR network at Day 14 ISO treatment. This is consistent with the human mutations preventing expression or function of these proteins to cause cardiac dysfunction, implying more maladaptive changes occurred at the prolonged phase. In contrast, Xin actin-binding repeat-containing protein 1 (XIRP1) was observed within the acute phase of ISO treatment. XIRP1 has been reported to be decreased upon acute cardiac stress as a protective mechanism (Otten, van der Ven et al. 2010, Wang, Lin et al. 2014). TTN was unique in this dataset as it was significantly up-regulated at the acute timepoint and significantly down-regulated at the prolonged timepoint. This could represent a possible adaptive up-regulation of titin acutely induced by stress and possibly representing hypertrophy preceding the maladaptive decrease in titin function observed in inherited cardiac diseases. Overall, based on the protein synthesis profile, it appears there may be an early cardioprotective phase followed by a maladaptive phase in MCR.

As we combined the NSP data with protein turnover rate measurements from in vivo ISO-treated mice compared to control mice, we identified a set of proteins falling into 4 distinct biological processes: Phagosome Maturation & Integrin-Linked Kinase Signaling, Oxidative Phosphorylation & Mitochondrial Function, Inflammatory Responses and Fatty Acid β-oxidation (Figures 4A **and** 4B). Proteins involved in phagosome maturation and ILK signaling were significantly enriched, exhibiting increased NSP levels in parallel with protein turnover rate, indicating that these biological processes are up-regulated during ISO-induced MCR. Phagosome maturation involves the activation of macrophages and neutrophils to degrade pathogens and cellular waste products in cardiomyocytes. Phagosomes degrade senescent cells and apoptotic cells to maintain cell homeostasis, and these groups of cells are affected during cardiac remodeling. In addition, ILK signaling is an essential process in the regulation of cardiac growth and hypertrophy. ILK is a multifunctional protein that physically links β-integrins with the contractile protein actin to regulate mechanoreceptors (Lu, Fedak et al. 2006, Nishimura, Kumsta et al. 2014). In our data, we observed an increase of NSP, and we observed that the rate of protein synthesis was greater than that of protein degradation (increased protein turnover, scenario 1 in **Table 1**) for proteins involved in phagosome maturation and ILK signaling, which falls in line with higher activities of these biological processes during ISO-induced MCR compared to sham treatment. These findings imply that apoptosis as well as the activation of macrophages and neutrophils in cardiomyocytes are important factors within MCR as previously reported and merit further study. (Narula, Haider et al. 1996, Olivetti, Abbi et al. 1997).

**Table 1:** Integration of protein turnover and PALM datasets for 14 days of ISO treatment. The table listed the four scenarios observed with this analysis and our proposed underlying mechanisms.

| <u>Scenario</u> | <u>NSP</u> | <u>Turnover</u> | <u>Proposed Mechanism</u> |
| --- | --- | --- | --- |
| 1 | Increased | Increased | Increased translation |
| 2 | Decreased | Increased | Increased degradation |
| 3 | Increased | Decreased | Decreased degradation |
| 4 | Decreased | Decreased | Decreased translation |

Our data also reveal that the majority (50%) of the protein dynamics detected is relevant to the biological processes of oxidative phosphorylation and mitochondrial function (Figures 4A **and** 4B); these proteins exhibited decreased levels of NSPs in parallel to a rate of protein synthesis greater than that of protein degradation (increased protein turnover rate, scenario 2 in **Table 1**). Perturbations in mitochondrial homeostasis and metabolism are essential mechanisms in the pathophysiology of cardiac remodeling and HF (Chaanine, Sreekumaran Nair et al. 2017). Previous observations suggest that mitochondrial biogenesis, signaling, and oxidative capacity are preserved or increased in the early stages of cardiac remodeling in order to match the energy demand imposed by a detrimental stressor such as ISO. Subsequently, the transition to MCR and HF corresponds to a decrease in mitochondrial biogenesis and oxidative metabolism (Chaanine, Sreekumaran Nair et al. 2017). In our data, we observe a decreased level of NSPs involved in oxidative phosphorylation, but these proteins exhibited an increased rate of protein turnover. A rate of protein synthesis that is higher than the degradation rate while the level of NSPs is declining suggests an imbalance of mitochondrial homeostasis, with many dysfunctional proteins as well as impaired oxidative capacity in the face of high energy. These data fall in line with observations in HF patients and post-infarct mice with cardiac remodeling (Murphy, Ardehali et al. 2016, Chaanine, Sreekumaran Nair et al. 2017). Mitochondrial proteins may have a higher synthesis than degradation rate in MCR (scenario 2 in **Table 1**); however, both the synthesis and degradation rates are most likely much lower compared to the sham group, clarifying the lower levels of NSPs. More specifically, most components of the respiratory chain, including electron transport chain (ETC) complex I (NDUFA7, NDUFA12, NDUFA13, NDUFS1, NDUFS2, NDUFS7, and NDUFV1) and complex V (ATP1A1, ATP1A2, ATP5A1, ATP5B, ATP5C1, ATP5F1, ATP5H, ATP5J, and ATP5O) exhibited this trend. Complex I – IV expression levels and activity appear to be decreased in HF patients (PMID: 27126807), falling in line with our data that shows a decrease in mitochondrial NSPs (**Figure 4A**, scenario 2 in **Table 1**).

Since the initial observation by Levine et al., an extensive amount of information has been accumulating regarding the role of inflammation in the initiation and progression of HF due to ischemic injury, myocardititis, and long standing hypertension (Levine, Kalman et al. 1990). Many circulating pro-inflammatory cytokines such as TNF, IL-1, and Il-6 have been implicated in the process of MCR and HF (Dick and Epelman 2016). The inflammatory response in MCR is closely associated with the activation of the immune system. Notably, cardiomyocytes are a significant source of pro-inflammatory mediators that contribute to the progression of MCR and HF. Our data confirm that proteins in the cardioprotective LXR activation pathway and inflammation pathways were enriched due to decreased degradation and a corresponding increase in NSPs (decreased turnover rate, scenario 3 in **Table 1**). Accordingly, inflammatory proteins may have a lower synthesis than degradation rate in MCR; however, both the synthesis and degradation rates are most likely higher compared to the sham group, clarifying the higher levels of NSPs. Thus, these data suggest that inflammatory processes are increased in ISO-induced MCR compared to sham treatment.

Interestingly, fatty acid oxidation has been reported to directly correlate with mitochondrial density and oxidative capacity, and a failing heart exhibits a metabolic shift towards glucose oxidation (Noordali, Loudon et al. 2017). Our data reveal that NSP levels in fatty acid β-oxidation are reduced while the protein degradation rate is higher than its synthesis, confirming that fatty acid oxidation is diminished during ISO-induced cardiac remodeling compared to sham treatment (**Figure 4A**, scenario 4 in **Table 1**). This observation also corresponds to the reported reduction in the protein expression of this pathway in cardiac remodeling as the fuel source shifts from fatty acids to carbohydrates (Akhmedov, Rybin et al. 2015).

In conclusion, the balance between protein synthesis and degradation is the basis for homeostasis for any proteome, but the paucity of cellular proliferation in the heart increases the importance of this delicate balance in its response to stress. MCR is a dynamic process that first attempts to restore homeostasis that was disrupted by physiological stressors but eventually produces deleterious changes to the proteome after prolonged stress, leading to cardiac dysfunction. Replicating known protein alterations in the cardiac literature provides confidence in the novel protein dynamics in our MCR network, which will provide deeper insights into cardiac human physiology. The identification and timing of adaptive and maladaptive changes induced by MCR is poorly understood. As our dataset provides a step to deconvolute the temporal dynamics of MCR, future functional studies will be required to confidently assign adaptive/maladaptive labels to the proteins in our MCR network. Our combination of NSP and turnover analysis suggests that the drugs that specifically manipulate translation and degradation (i.e., proteasomal or autophagic) processes could alter biological pathways differentially and possibly produce different clinical outcomes. Finally, we propose that the novel integration of NSP and turnover proteomic datasets could be applied to all animal models of disease to provide a greater understanding of the dysregulation of homeostasis in human diseases.

## Acknowledgements

The study was supported by NIH grants U54GM114833 (to P.P and J.R.Y), R35HL135772 (to P.P) and P41GM103533 (to J.R.Y.)

## Supplementary Legend

Supplementary Table 1: The number of identified newly synthesized proteins and peptides for each MS analysis.

Supplementary Table 2: The NSPs that were significantly (p<0.05) altered in the Day 4 ISO analysis.

Supplementary Table 3: The NSPs that were significantly (p<0.05) altered in the Day 14 ISO analysis.

Supplementary Table 4: Large unquantifiable NSP changes in the Day 4 ISO analysis.

Supplementary Table 5: Large unquantifiable NSP changes in the Day 14 ISO analysis.

Supplementary Table 6: NSPs that were statistically significant with ISO at both the Day 4 and Day 14 analyses.

Supplementary Table 7: Significantly altered NSPs in this study previously associated with cardiac dysfunction in the literature.

Supplementary Table 8: Quantified NSPs compared to their previously reported turnover rates.

---

**Supplementary Table 1:** Identifications. H-ISO/L-Sham means this replicate is proteins mixed from H-AHA-labeled ISO mice and L-AHA-labeled sham mice in a 1:1 ratio after protein assay. H-sham/L-ISO was named in the same way. All proteins were required to be identified with at least 1 peptide. All peptides were required to be fully tryptic. For all datasets, both protein FDR and peptide FDR were <1%.

| Labeling Time frame | Replicate | Content | AHA Protein | AHA Peptide |
| --- | --- | --- | --- | --- |
| Day 1-4 | A | H-ISO/L-Sham | 2,189 | 9,522 |
|  | B | H-ISO/L-Sham | 1,515 | 5,493 |
|  | C | H-sham/L-ISO | 1,232 | 3,811 |
| Day 10-14 | E | H-ISO/L-Sham | 1,053 | 3,581 |
|  | F | H-ISO/L-Sham | 1,306 | 5,511 |
|  | G | H-sham/L-ISO | 1,574 | 6,899 |

